# Diversity and evolution of chromatin regulatory states across eukaryotes

**DOI:** 10.1101/2025.03.17.643675

**Authors:** Cristina Navarrete, Sean A. Montgomery, Julen Mendieta, David Lara-Astiaso, Arnau Sebé-Pedrós

**Affiliations:** Centre for Genomic Regulation (CRG), Barcelona Institute of Science and Technology (BIST), Barcelona, Spain; Universitat Pompeu Fabra (UPF), Barcelona, Spain; Clinica Universidad de Navarra, Centro de Investigacion Medica Aplicada (CIMA), Instituto de Investigación Sanitaria de Navarra (IDISNA), Pamplona, Spain; Arc Institute, Palo Alto, CA, USA; ICREA, Barcelona, Spain; Tree of Life Program, Wellcome Sanger Institute, UK

## Abstract

Histone post-translational modifications (hPTMs) are key regulators of chromatin states^1,2^, influencing gene expression, epigenetic memory, and transposable element repression across eukaryotic genomes. While many hPTMs are evolutionarily conserved^3^, the extent to which the chromatin states they define are similarly preserved remains unclear. Here, we developed a combinatorial indexing ChIP-seq method to simultaneously profile specific hPTMs across diverse eukaryotic lineages^4^, including amoebozoans, rhizarians, discobans, and cryptomonads. Our analyses revealed highly conserved euchromatin states at active gene promoters and gene bodies. In contrast, we observed diverse configurations of repressive heterochromatin states associated with silenced genes and transposable elements, characterized by various combinations of hPTMs such as H3K9me3, H3K27me3 and/or different H3K79 methylations. These findings suggest that while core hPTMs are ancient and broadly conserved, their functional readout has diversified throughout eukaryotic evolution, shaping lineage-specific chromatin landscapes.

## Main

Access to genetic information in eukaryotes is regulated by a nucleoproteic interface known as chromatin. Nucleosomal chromatin establishes a repressive ground state for transcription and other DNA-templated processes in eukaryotic genomes^5,6^. The regulatory environment and activity of different genomic regions are determined by distinct combinations of chemical modifications^7–11^, chromatin components^12,13^, and structural features^14^. These functional configurations, often referred to as chromatin states, change dynamically during cell differentiation^15,16^ and can be inherited across cell divisions^17^. Furthermore, chromatin-based regulation plays a crucial role in repressing parasitic genomic elements, such as transposable elements (TEs) and viruses^18–22^.

Post-translational modifications of histone tails (hPTMs) are central to defining functional chromatin states^1,23–25^. These hPTMs are conserved across diverse eukaryotes^26–29^, with dozens tracing back to the last eukaryotic common ancestor (LECA)^3^. In this study, we investigated the diversity of chromatin states defined by these conserved hPTM across major eukaryotic lineages (**Fig. 1a**). To this end, we developed iChIP2, a versatile and low-input combinatorial ChIP-seq method that enables large scale hPTM analysis by pooling indexed chromatin from diverse species into a single chromatin immunoprecipitation reaction^15^ (**Fig. 1b**). Using iChIP2, we efficiently tested 25 anti-hPTM antibodies across multiple species simultaneously and selected the most optimal for each hPTM (**Extended Data Fig. 1**). Then, we profiled a total of 12 hPTMs across 12 phylogenetically diverse species that collectively represent major eukaryotic lineages (**Fig. 2, Extended Data Fig. 2**). We combined these genome-wide maps with gene expression data and various genomic features (e.g. TEs and promoters) to functionally interpret chromatin states.

**Figure 1.**
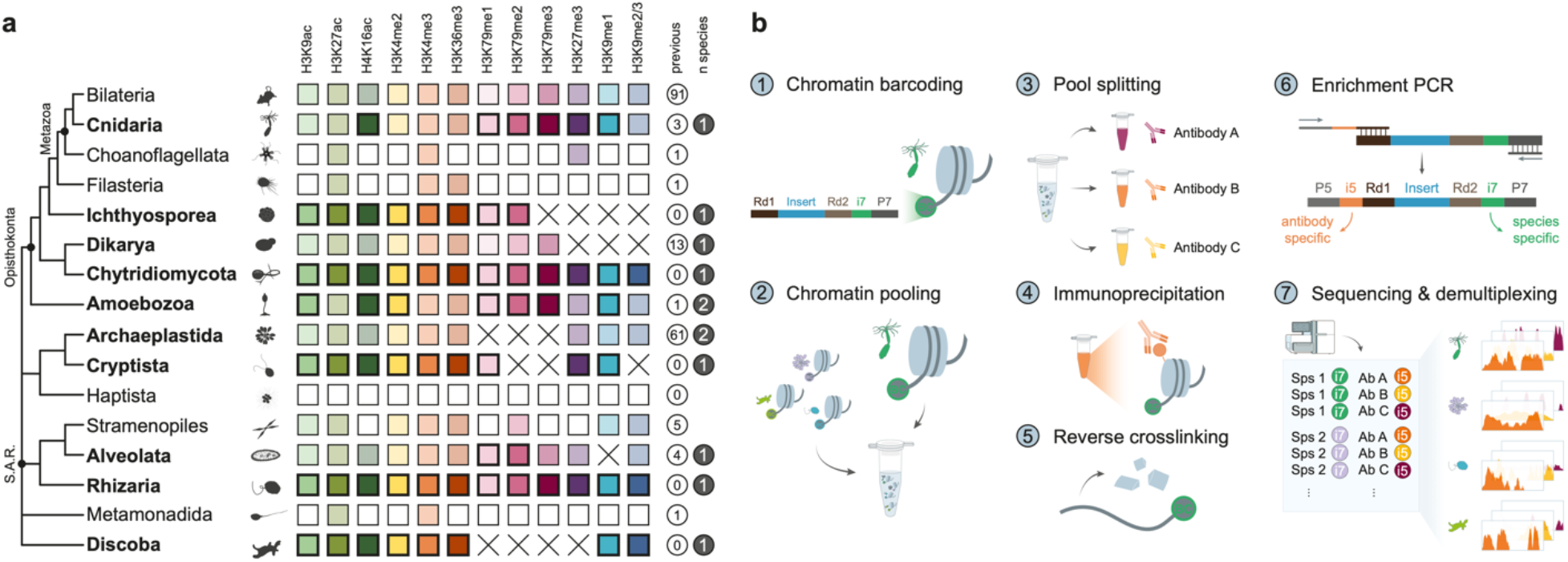
iChIP2 enables multiplexed chromatin profiling across eukaryotes. **a**, Eukaryotic tree of life, highlighting the lineages examined in this study. Highlighted, colored boxes represent newly profiled histone modifications in this study, while semi-transparent, colored and white boxes denote whether data for a particular histone modification was available or unavailable within a lineage, respectively. Crosses denote the absence of a histone modification in certain eukaryotic lineages. Numbers inside the white circles indicate the number of species with histone modification ChIP-seq data in GEO for each branch, while the numbers inside dark circles specify the number of species profiled per lineage in this study. **b**, Schematic illustration of the iChIP2 protocol. The iChIP2 involves initial sample-specific chromatin barcoding, followed by pooled chromatin immunoprecipitation using target antibodies. The last step adds a second index to identify each antibody after sequencing.

**Figure 2.**
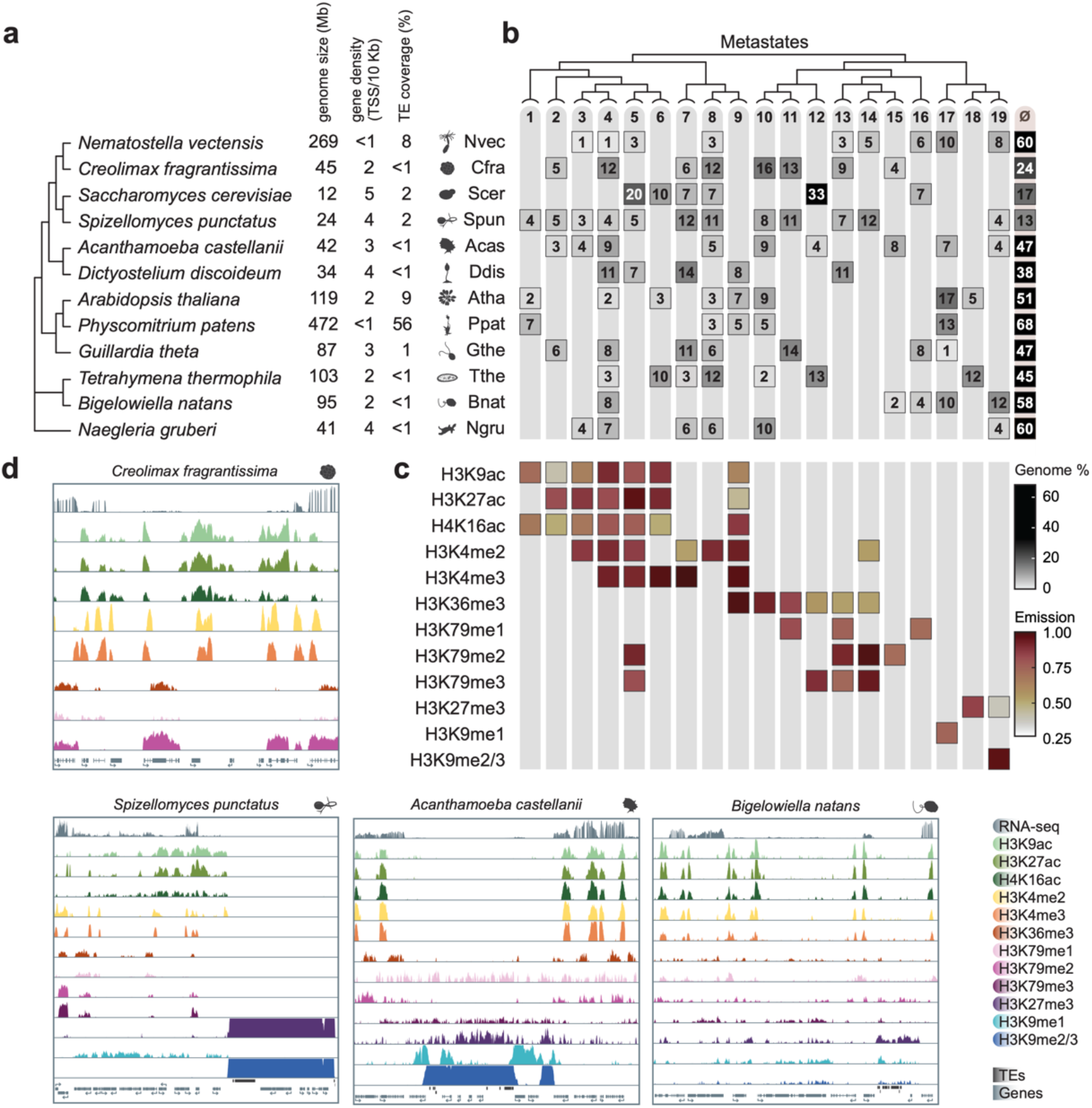
Cross-species chromatin states. **a**, Species tree sampled for iChIP2 experiments. Genome size (Mb), gene density (average number of transcription start sites per 10kb, and the percentage of the genome covered by transposable elements are plotted beside each species name. **b**, Heatmap of the percentage of the genome of each species corresponding to each metastate. The simplified hierarchical clustering tree is displayed above the heatmap. **c**, Contribution of each profiled histone modification to each metastate. Values are calculated as the median emission probabilities of all ChromHMM states grouped into one metastate. **d**, Browser snapshots of profiled histone modifications. RNA-seq reads (grey, top), histone modifications, transposable elements (black boxes, bottom), and genes (grey, bottom) are shown.

To integrate genome-wide hPTM profiling data across all species, we first defined chromatin states using ChromHMM^30^ for each species and manually selected models that best captured chromatin state diversity while minimizing redundancy. We then clustered chromatin states across species by comparing the emission probabilities of shared hPTMs (**Extended Data Fig. 3**), revealing three broad cross-species clusters: (1) states proximal to transcription start sites (TSS), (2) active gene bodies, and (3) repressed genomic features, including TEs and silent genes. To further refine these clusters, we collapsed similar chromatin states into broader categories, which we term metastates (**Fig. 2b-c, Extended Data Fig. 3**). The final 19 metastates represented distinct combinations of histone modifications. Among them, metastates 1-9 were dominated by histone acetylation and H3K4me2/3 around promoters, metastates 10-16 comprised primarily H3K36me3 and H3K79me1/2/3 over gene bodies, and metastates 17-19 were enriched in heterochromatic hPTMs like H3K9me2/3 or H3K27me3 (**Fig. 2c, Extended Data Fig. 3**). No single metastate was present in all species; however, metastates 4 and 8 were observed in all but two species (**Fig. 2c**). While metastate 4 encompassed all profiled histone acetylation marks plus H3K4me2/3, metastate 8 was defined solely by H3K4me2 (**Fig. 2b-c**). Additionally, we calculated the percentage of the genome in each species associated with each metastate (**Fig. 2b**). In all species, a substantial fraction of the genome (13–68%) was not assigned to any chromatin state (null state). Species with small, gene-dense genomes such as *C. fragrantissima* or *G. theta* were predominantly characterized by active chromatin states enriched in histone acetylation (metastates 1-6), as well as active gene body stated marked by H3K36me3 (metastates 9-12). In contrast, species with larger genomes, such as *N. vectensis, B. natans* or *P. patens*, contained a significant fraction of their genome (13-28%) dominated by repressive hPTMs (metastates 17-19).

To investigate the evolution of chromatin state and their specific functions across eukaryotes, we analyzed the relationship between gene expression and histone modifications at promoter regions and gene bodies. To this end, we clustered genes in each species based on hPTM coverage across 10bp bins covering the gene body and then sorted these clusters based on RNA-seq expression levels (**Extended Data Fig. 4**). We examined the distribution of hPTMs around the TSS in gene clusters representing expressed and non-expressed genes (**Fig. 3a**). In all species, expressed genes exhibited high H3K4me2/3 and acetylation signal downstream of the TSS, followed by high H3K36me3 levels over the gene body (**Fig. 3a**), a configuration previously observed in various eukaryotes^8,16,29,31^. This confirms H3K4 methylation and histone acetylation as universal activation marks. Moreover, we systematically searched for a characteristic bimodal H3K4me3 distribution is often found around the TSS in some animal species and that has been linked to anti-sense transcription at these sites^8,16^. We controlled for artifacts (see Methods) such as close head-to-head genes or unannotated genes revealed by RNA-seq (**Extended Data Fig. 5**), confirming that this bimodal H3K4me3 pattern is unique to bilaterian animals.

**Figure 3.**
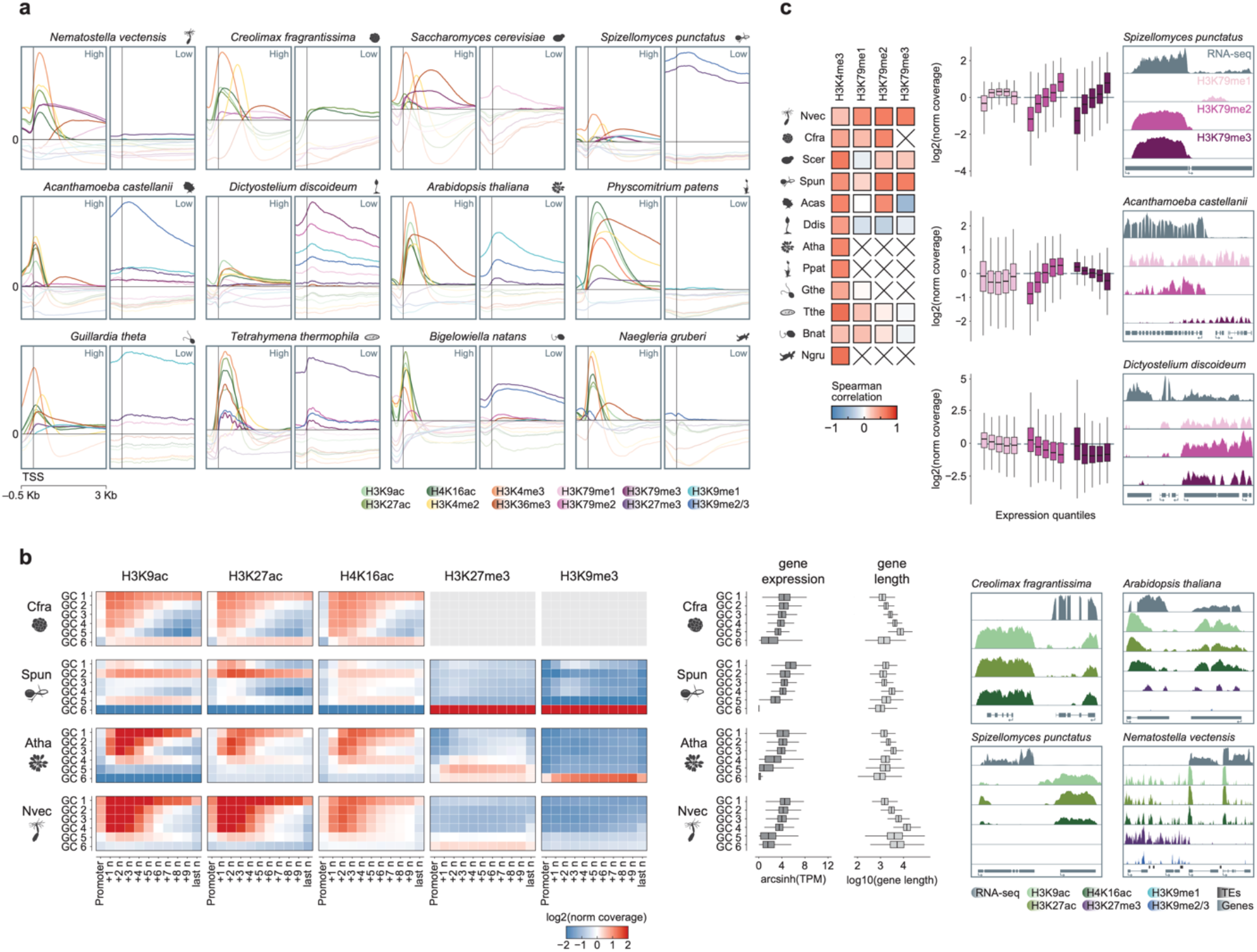
Chromatin states and gene expression. **a**, Distribution of histone modifications within active and repressed genes. Profile plots show average read coverage of a histone modification, log2 normalized to input. Profiles extend 500bp before and 3kb after the transcription start site (TSS). Solid vertical lines indicate the TSS. Solid horizontal lines indicate a value of 0, with values below zero displayed as transparent lines. Genes are selected from k-means clusters calculated in Extended Data Fig. 4. **b**, Heatmaps of histone mark enrichment per k-means gene cluster over genes. Genes were divided into 200bp bins, representing roughly one nucleosome, and the average read coverage of the indicated histone modification was calculated. Genes shorter than the indicated bin were not included in the average values of that bin. Promoters include 500bp upstream of the TSS and the last nucleosome (last n) comprises the last 200bp of genes longer than 1.8kb. Gene clusters are ordered by average gene expression, shown in boxplots to the right of the heatmaps. Gene lengths per gene cluster are also shown in boxplots. Browser snapshot tracks follow the same colour scale as profile plots above, in addition to RNA-seq reads (grey, top) transposable elements (black boxes, bottom), and genes (grey, bottom). **c**, Spearman correlation of average histone modification coverage, log2 normalized to input, with gene expression. Crossed out boxed indicate species lacking the histone modification. Boxplots show average histone modification coverage for H3K79me across gene expression quintiles, ordered left to right from lowest to highest gene expression. Browser snapshots follow the same colour scale as boxplots, in addition to RNA-seq reads (grey, top) and genes (grey, bottom).

In some species, we observed an unexpected association between histone acetylation and the gene bodies of lowly expressed (but not fully repressed) genes (**Extended Data Fig. 4**). Even after controlling for gene length effects, this pattern of broad gene body acetylation in lowly expressed genes was evident in *C. fragrantissima* (GC6), *S. punctatus* (GC5) and in *A. thaliana* (GC4) (**Fig. 3b**), while this was absent in other species, such as *N. vectensis* (**Fig. 3b**). In all three species, gene body histone acetylation occurred in the absence of post-TSS H3K4me2/3 signal (**Extended Data Fig. 4**). Additionally, in *S. punctatus* and *A. thaliana*, this acetylation pattern was observed outside the context of heterochromatin (H3K27me3 and/or H3K9me2/3). Of note, a similar pattern was previously reported in the filasterean *Capsaspora owczarzaki*^*16*^. Gene body histone acetylation has been linked to high levels of expression and BRD4 occupancy in vertebrates^32^, but the specific writer/reader mechanisms responsible for gene body acetylation in lowly expressed genes in chytrid fungi, plants, and ichthyosporeans remain unclear and warrant further investigation.

Our analysis also revealed distinct associations between H3K79 methylation and gene expression levels (**Fig. 3a, Extended Data Fig. 4**). To systematically quantify these effects, we calculated the correlation between gene expression levels and H3K79me1/2/3 coverage over gene bodies. In most species, we observe a positive correlation between H3K79me1/2/3 levels and expression levels (**Fig. 3c**), similar to what is observed in vertebrates^33^. However, we identified multiple exceptions to this pattern. For example, gene expression was negatively correlated with H3K79me1 coverage in *S. cerevisiae*, and with H3K79me3 coverage in *T. thermophila* and *B. natans* (**Fig. 3c**). The most striking pattern was observed in amoebozoans. In *D. discoideum* all three H3K79 methylations were negatively associated with gene expression, whereas in *A. castellani*, each methylation state displayed distinct correlations: none for H3K79me1, positive for H3K79me2, and negative for H3K79me3 (**Fig. 3c**). In animals and yeast, H3K79 methylation is catalyzed by the phylogenetically conserved Dot1/DOT1L methyltransferase family^33^. While the Dot1 protein in most species lacks accessory domains, one *A. castellanii* Dot1 homolog contains a chromo domain, often involved in binding methylated histone residues, and another homolog possesses an MBD domain, known to be involved in DNA methylation binding (**Extended Data Fig. 5**). We hypothesize that these accessory domains could target these Dot1 homologs to repressed regions marked with H3K9me3 (see below) and DNA methylation. Additionally, a third Dot1 homolog in *A. castellanii* contains a phenylalanine residue in the H3K79-binding channel, a feature previously identified in the trypanosome *T. brucei* and known to restrict the conversion of H3K79me2 to H3K79me3^34^. This *A. castellanii* Dot1 homolog, which lacks accessory domains, could be responsible for depositing H3K79me2 over transcribed gene bodies. Overall, H3K79 methylation exemplifies the phylogenetic diversity in the relationship between hPTMs and gene expression, highlighting its evolutionary variability across lineages.

Having revealed this unexpected link between H3K79 methylation and repressive states, we examined in depth the diversity of heterochromatic states across eukaryotes, focusing both on non-expressed genes (**Extended Data Fig. 4**) and on TEs. To ensure consistency, we first systematically reannotated TEs in each species^35^. We then clustered TEs based on hPTM coverage (**Extended Data. Fig. 6**) and associated each cluster to major TE classes (**Extended Data Fig. 7**). By analyzing the combinations of repressive hPTMs and their associated genomic features, we defined distinct heterochromatic states in each species (**Fig. 4a, Extended Data Fig. 8**). In the non-bilaterian animal *N. vectensis*, we identified H3K27me3 enriched over repressed genes (**Fig. 4b, Extended Data Fig. 4**) and two additional heterochromatic states associated with TEs: H3K9me3 was linked to retrotransposons (LINEs, SINEs, LTRs), while a combination H3K27me3 and H3K9me1 was associated with TIR and Helitron DNA TEs (**Fig. 4b, Extended Data Figs. 6, 7 and 8)**. These findings suggest that a dual role for heterochromatin in repressing gene expression and TEs is phylogenetically widespread across metazoa^8^, including non-bilaterians. Among fungi, many species such as *S. cerevisiae* have lost all known repressive hPTMs, but different heterochromatic states are present in others^36–38^. Here, we analyzed heterochromatin in an early-branching fungal lineage, represented by the chytrid *S. punctatus*, and found that both TEs and non-expressed genes were strongly associated with high H3K27me3 and H3K9me3 levels (**Fig. 4b, Extended Data Fig 6 and 7**). In comparison, H3K9me3 and H3K27me3 were mutually exclusive over genes but not TEs in the ascomycete *Podospora anserina*^36^, while both marks are antagonistic in other species^37,38^, further highlighting the dynamic evolution of heterochromatin within fungi. In plants we also identified different heterochromatic states (three in *A. thaliana*, similar to those previously described^39,40^; and two in the moss *P. patens*^41^), which are similar to those observed in other land plants^9^ and in unicellular green algae^42^, while in the ciliate *T. thermophila* we identified a single repressive state in the defined by H3K27me3 and corresponding to repressed genes in the macronuclear genome.

**Figure 4.**
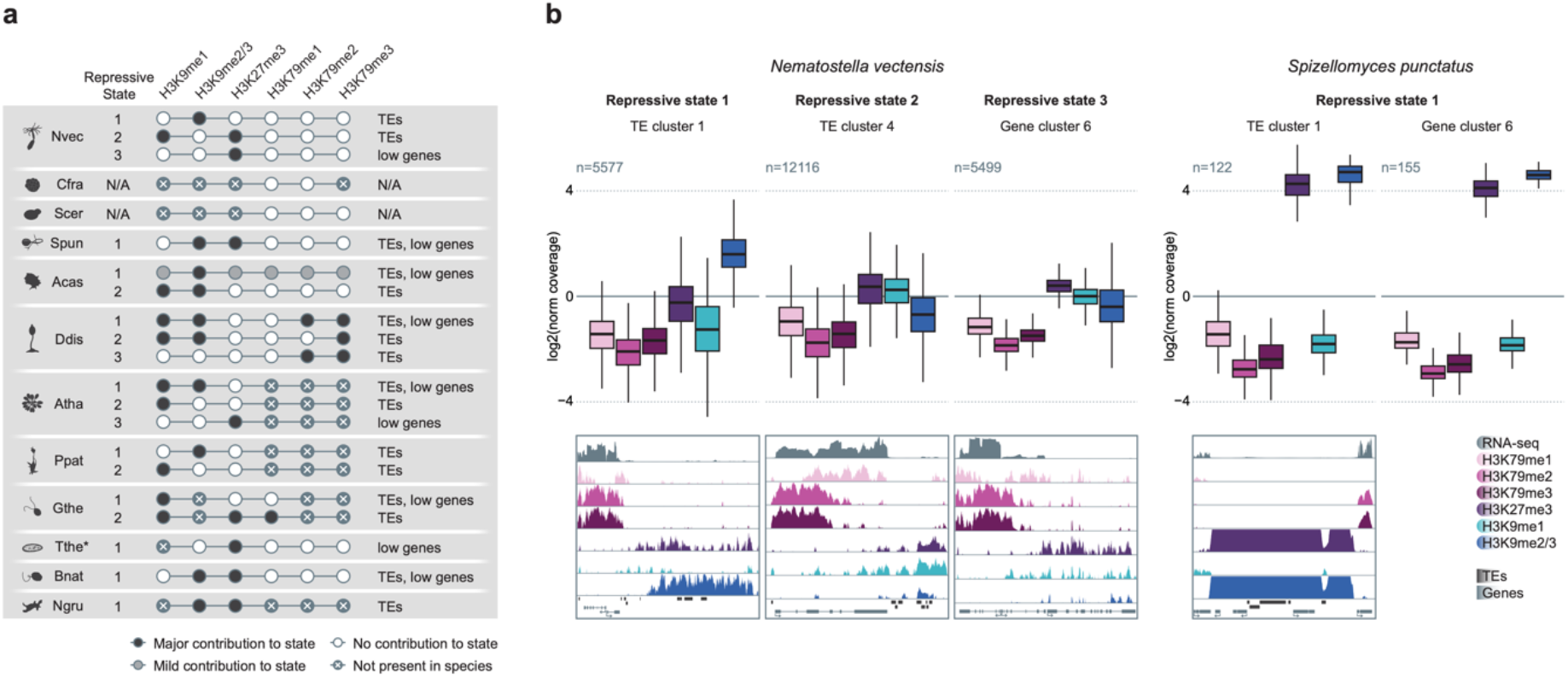
Diversification of repressive chromatin states. **a**, Summary table of distinct types of heterochromatin within each species. Black circles indicated a strong presence of the histone modification, grey circles indicate a weak presence of the histone modification, white circles indicate an absence of the histone modification in that repressive state, and white crosses on grey backgrounds indicate a complete absence of the histone modification in the species. Lines connecting circles indicate a co-occurrence of the histone modifications within the repressive state. Grey circles with crosses indicate absence of the histone modification in the species^3^. Genomic features associated with each type of heterochromatin are noted, TEs for transposable elements and low genes for lowly expressed genes. **b**, Boxplots of average read coverage of a histone modification, log2 normalized to input, of individual transposable elements within a selected cluster from Extended Data Fig. 6. Browser snapshots follow the same colour scale as boxplots, in addition to RNA-seq reads (grey, top), transposable elements (black, bottom), and genes (grey, bottom).

Repressive hPTMs had not been previously studied in amoebozoans. Here, we identified two distinct heterochromatin states in *A. castellani* (**Fig. 4a, Extended Data Fig. 8**): (1) H3K9me3 with weaker H3K27me3 and H3K79me1/3, associated with TEs and repressed genes, and (2) H3K9me3 with H3K9me1, linked to a separate subset of TEs. In *D. discoideum*, we identified three heterochromatin states (**Fig. 4a, Extended Data Fig. 8**): (1) H3K9me1/3 and H3K79me2/3, associated with both repressed genes and TEs, (2) a similar state lacking H3K79me2, and (3) a distinct combination of only H3K79me2/3, associated with different TE subsets. We also characterized previously unsampled eukaryotic lineages, including cryptomondas (*G. theta*), rhizarians (*B. natans*) and discobans (*N. gruberi*) (**Fig. 4a, Extended Data Fig. 8**). In *G. theta* we identified two heterochromatin states: (1) H3K9me1 associated with TEs and (2) a state enriched in H3K9me1, H3K27me3, and H3K79me1, linked to repressed genes and a distinct set of TEs. In *B. natans* and *N. gruberi*, we identified a single heterochromatic state defined by high H3K9me3 and H3K27me3, but with different genomic associations: in *N. gruberi*, this state was exclusively associated with TEs (but not repressed genes), whereas in *B. natans*, it was linked to both TEs and repressed genes. Overall, our analysis reveals a striking diversity of repressive states across eukaryotes, ranging from complete absence (*S. cerevisiae* and *C. fragrantissima*) to distinct hPTM associations with genomic features and transcriptional states, and adding to those that have been previously described in other lineages^43,44^. Interestingly, a similar diversity has been observed in DNA methylation patterns across eukaryotes^45,46^. We hypothesize that the evolutionary driver behind this diversity is the repression of TEs and other parasitic genomic elements, such as viruses. This evolutionary arms race between TEs and host repression mechanisms is exemplified by the repeated capture of chromatin reader domains by TEs^3,47^ or the acquisition of chromatin modifiers by endogenized viruses^48^. Over evolutionary timescales, these TE-targeting repressive mechanisms may have been co-opted for gene regulation in different lineages, leading to the broad diversity of gene expression repressive chromatin states observed across eukaryotes.

In summary, our combinatorial barcoding ChIP-seq method enables efficient, multiplexed profiling of hPTMs across diverse eukaryotic species. The results we obtained using iChIP2 reveal that, despite the broad conservation of many hPTMs across eukaryotes^3^, the functional chromatin states defined by these marks are not always conserved. This is exemplified by the changing relationship between H3K79me1/2/3 and gene expression levels, as well as the different repressive heterochromatin configurations associated to TEs. These findings may indicate evolutionary changes in reader proteins that link hPTMs to different chromatin-modifying catalytic activities, such as nucleosomal remodeling, hPTM writing/erasing, and/or DNA methylation. This hypothesis is supported by the rapid phylogenetic diversification of these reader proteins compared to the relatively conserved catalytic chromatin proteins (writers, erasers and remodellers)^3^. More broadly, our results also exemplify the potential of biodiversity epigenomic profiling. At the time when genome sequencing is rapidly advancing across the tree of life^49^, this approach offers a valuable opportunity to similarly expand our understanding of genome function and regulation.

## Author contributions

D.L. and A.S-P. conceived the project. C.N and D.L. performed experiments. S.A.M., C.N., J.M. and A.S-P. performed data analysis. S.A.M and C.N. generated data visualizations. D.L. and A.S-P. provided supervision. S.A.M, C.N., and A.S-P. wrote the manuscript with contributions from all authors.

## Acknowledgments

Research in A.S-P. group was supported by the European Research Council (ERC-StG 851647) and the Spanish Ministry of Science and Innovation (PID2021-124757NB-I00. C.N. is supported by FPI PhD fellowships from the Spanish Ministry of Science and Innovation. S.A.M. was supported by the European Union’s H2020 research and innovation programme under Marie Sklodowska-Curie grant agreement 101110098.

## Methods

### Culture Conditions

*Acanthamoeba castellanii* strain Neff (ATCC 30010) and *Naegleria gruberi* strain ATCC30224 were axenically grown in ATCC mediums 712 PYG with additives and 1034 (modified PYNFH medium) respectively at 23 °C. *Dictyostelium discoideum* strain AX4 was axenically grown in HL5-C with glucose medium at 23 °C, *Creolimax fragrantissima* strain CH2 was axenically grown in Marine Broth medium at 18 °C, *Spizellomyces punctatus* strain DAOM BR117 was axenically grown in a medium containing 0.5% yeast extract, 3% glycerol,1 g/l of K_2_HPO_4_ and 0.5% ethanol at 18 °C, *Tetrahymena thermophila* strain CU428 was axenically grown in SSP medium at 30 °C and maintained in an autoclaved vial with 10 mL MilliQ H_2_O and an organic chickpea at 23 °C and *Bigelowiella natans* strain CCMP2755 and *Guillardia theta* strain CCMP2712 were axenically grown in ATCC mediums L1-Si and Prov50, respectively, at 25 °C under a 12:12 h light-dark cycle. *Saccharomyces cerevisiae* strain BY4742 was grown in YPD medium in liquid cultures at 30 °C. *Nematostella vectensis* polyps were grown in 1/3X filtered artificial seawater at 18^°^C in the dark and fed with freshly hatched *Artemia salina* nauplii five times a week. Additionally, we examined *Arabidopsis thaliana* Col-0 accession leaf tissue and *Physcomitrium patens* Gransden accession gametophytes, provided courtesy of J. Casacuberta (Centre for Research in Agricultural Genomics-CSIC, Barcelona).

### Cell/tissue Crosslinking

Cells were harvested and washed in 1X PBS twice. Pellet was resuspended in 10 ml 1X PBS and 10 mL 2% methanol-free formaldehyde (ThermoFisher, 28906) was added (1:1 ratio, 1% final formaldehyde concentration). Cells were crosslinked for 10 min at room temperature (RT) with rotation. Crosslinking reaction was quenched with 0.75 M Tris-HCl pH 7.5 for 5 min at RT with rotation. Cells were pelleted and washed with 1X PBS. Crosslinked cell pellets were flash-frozen in liquid nitrogen and stored at −80 °C.

In the case of *Nematostella vectensis, Arabidopsis thaliana* and *Physcomitrium patens*, steps were similar but the crosslinking reaction was performed under vacuum in a borosilicate desiccator (VWR, 467-0086) for 10 min at RT with no rotation.

### Chromatin extraction and fragmentation

For all the species except plants, 5−10M crosslinked cells (or 2-3 crosslinked juvenile polyps in case of *N. vectensis*) were resuspended in 500 μl of Cell Lysis Buffer (20 mM HEPES pH 7.5, 10 mM NaCl, 0.2% IGEPAL CA-630, 5 mM EDTA) supplemented with 1X protease inhibitors cocktail (PIC, Roche, 4693132001), and incubated on ice for 10 min. Cells were centrifuged at max speed for 10 min at 4 °C, and the resulting pellets were resuspended in 100 µl Bead Beating Buffer (20 mM HEPES pH 7.5, 10 mM NaCl, 5 mM EDTA) + 1X PIC. Samples were transferred to 0.2 ml tubes containing 1/3 acid-washed glass beads (G8772, Sigma-Aldrich) and vortexed for 30” five times with 30” incubation on ice between each. The resulting lysate was moved to a 1.5 ml Bioruptor microtube (Diagenode, C30010016), and sodium dodecyl sulfate (SDS, Sigma-Aldrich, 71736) was added to 0.6%. Chromatin was sheared for 5−8 cycles of 30” ON / 30” OFF using a Bioruptor Pico (Diagenode) in order to generate 150−300 bp fragments. To reduce the SDS concentration, chromatin was diluted with five volumes of Dilution Buffer (20 mM HEPES pH 7.5, 140 mM NaCl), and centrifuged at max speed for 10 min at 4 °C. The supernatant containing solubilized, sheared chromatin was aliquoted and stored at −80 °C, except for 60 µl (10%) used to QC the chromatin prep.

In the case of plants, chromatin extraction was performed as previously described (Payá-Milans et al., 2019) with minor modifications. Briefly, ~0.5 mg of plant tissue was ground with a mortar a pestle in liquid nitrogen to a fine powder. After adding 10 ml of Extraction Buffer 1 (0.4 M sucrose, 10 mM Tris-HCl pH 8, 10 mM MgCl_2_, 5 mM β-mercaptoethanol) + 1X PIC, cell suspension was filtered using a 70 µm cell strainer (pluriSelect Life Science, 43-50070-51) to clear any large tissue fragments and centrifuged at 5.000x*g* for 10 min at 4 °C. The cell pellet was resuspended in 1 ml Extraction Buffer 2 (0.25M sucrose, 10 mM Tris-HCl pH 8, 10 mM MgCl_2_, 1% Triton X-100, 5 mM β-mercaptoethanol) + 1X PIC, and centrifuged at 5000x*g* for 10 min at 4 °C. In order to purify the nuclear fraction, the pellet was resuspended in 300 µl Extraction Buffer 3 (1.7M sucrose, 10 mM Tris-HCl pH 8, 2 mM MgCl_2_, 0.15% Triton X-100, 5 mM β-mercaptoethanol) + 1X PIC, placed over 600 µl Extraction Buffer 3 in a new 2 mL tube and centrifuged at 16.000x*g* for 1h at 4 °C. Nuclei were resuspended in 200 µl Nuclei Lysis Buffer (50 mM Tris-HCl pH 8, 10 mM EDTA, 1% SDS) + 1X PIC and transferred to a 1.5 ml Bioruptor microtube. Chromatin was sonicated for 20−30 cycles of 30” ON / 30” OFF using a Bioruptor Pico (Diagenode). Chromatin was diluted 10 times adding ChIP Dilution Buffer (16.7 mM Tris-HCl pH 8, 1.2 mM EDTA, 167 mM NaCl, 1.1% Triton X-100), and centrifuged at max speed for 10 min at 4^°^C. Supernatant was stored at −80 °C in aliquots.

### iChIP2

We developed a multiplexed, low input ChIP-seq method, that include the following main steps (a detailed protocol is available in **Supplementary Note 1**):

1. Chromatin barcoding
2. Chromatin samples pooling
3. Immunoprecipitation
4. Stringent washes and reverse crosslinking
5. Library amplification

The initial sample-specific indexing (i7) step enables pooling barcoded chromatin from different samples, thereby reducing the individual chromatin amount per sample. A minimum of 100 ng (+10%) of chromatin per sample is recommended, and the use of fewer than 5 samples at this scale is not advised. Each pool is then divided into two or three distinct immunoprecipitation reactions, depending on the utilized antibodies, and incubated overnight. Following washings and reverse crosslinking steps, the target DNA is amplified via PCR and an antibody-specific index (i5) is added resulting in the final ChIP library.

We regularly profile 8 histone modifications from 8 different species at a time, generating 64 libraries in a single experiment. Both pre- and post-chromatin immunoprecipitation steps are conducted in 96-well plates, making our protocol scalable and suitable for automation on a liquid handling platform.

#### Chromatin Barcoding

To adjust the buffer for the adaptors ligation reaction, sheared chromatin was diluted 10 times by adding EB buffer (10 mM Tris-HCl pH 8) + 1X PIC, transferred into a 0.5 ml Amicon (30 KDa, Millipore, UFC503096) and centrifuged at 10.000x*g* for 10 min at 4 °C. Amicon filter was turned upside down onto a new 1.5 ml DNA low binding tube and spun down for 2 min at 100x*g* at 4 °C to collect the concentrated chromatin sample, transferred to a 96-well plate and the volume was brought to 50 µl with EB buffer + 1X PIC. Chromatin End Repair and A-tailing was performed by adding 3 µl NEB Next Ultra II End Prep Enzyme Mix + 7 µl NEB Next Ultra II End Prep Reaction (NEB, E7645L). Samples were incubated in a thermocycler at 20 °C for 45 min, 50 °C for 20 min and cooled to 4 °C. After end repair, chromatin was indexed by adding 5 µl of 2.5 mM custom Y-shapped i7 adaptors (**Supplementary Table 1**) and 30 µl NEB Next Ultra II Ligation Master Mix + 1 µl NEB Next Ultra II Ligation Enhancer (NEB, E7645L) to each well. Samples were thoroughly mixed, spun down and incubated in a thermal cycler at 20 °C for 45 min, 16 °C for 45 min, 12 °C for 45 min and cooled to 4 °C. The barcoding capacity of 5 µl 2.5 mM custom adaptors is up to 400 ng (+10%) of chromatin.

#### Chromatin Pooling

Barcoded chromatin samples from different species were combined by pipetting each into the same 15 ml Amicon filter (50 KDa, Millipore, UFC905024) containing 10 ml Centricon buffer (10 mM Tris-HCl pH 8, 140 mM NaCl, 1 mM EDTA, 0.1% SDS) + 1X PIC. This removed any residual adaptors and concentrated the final pool in a buffer suitable for the immunoprecipitation step. To recover as much chromatin as possible, each well was washed with 200 µl Centricon buffer and transferred to the 15 ml Amicon filter. The Amicon filter was mixed by inversion 10-12 times and spun down at 2000x*g* for 20 min. The concentrated volume (~200 µl) was transferred to a 1.5 ml protein low binding tube and Triton X-100 was added to 1% final concentration. Pooled chromatin samples were brought to 550 µl with RIPA buffer (10 mM Tris-HCl pH 8, 140 mM NaCl, 1 mM EDTA, 0.1% SDS, 1% Triton X-100) + 1X PIC, and divided into two protein low binding tubes with 250 µl each for two independent chromatin immunoprecipitations. The remaining 50 µl were kept as barcoded input.

#### Immunoprecipitation

Antibody was added to a pooled indexed chromatin sample and incubated at 4^°^C for 12−14 h with rotation. For each ChIP, we used 5 µg anti-H3K9ac (Millipore, 17-658), 2.5 µg anti-H3K27ac (Abcam, ab4729), 3 µg anti-H3K4me2 (Abcam, ab32356), 2.5 µl anti-H3K4me3 (Millipore, 07-473), 3 µg anti-H3K36me3 (Abcam, ab9050), 8 µg anti-H3K79me1 (Abcam, ab177183), 4 µg anti-H3K79me2 (Abcam, ab3594), 2.5 µg anti-H3K79me3 (Diagenode, C15410068), 5 µl anti-H4K16ac (Millipore, 07-329), 2 µg anti-H3K9me1 (Abcam, ab9045), 3 µg anti-H3K9me3 (Abcam, ab176916), 8 µg anti-H3K27me3 (Millipore, 07-449) and 2 µg anti-H3 (Abcam, ab1791). After O/N incubation, 10 µl Protein A (Millipore, 16-661) + 10 µl Protein G (Millipore, 16-662) magnetic beads were washed twice with RIPA buffer, resuspended in 25 µl RIPA buffer + 1X PIC, added to the IP samples and incubated at 4^°^C for 2 h with rotation. An overview of the antibodies tested across different species is available in **Extended Data Fig. 1**.

#### Washes and reverse crosslinking

IP samples were magnetized, supernatant was removed, and beads were resuspended in 175 µl RIPA buffer + 1X PIC and transferred to a 96-well plate. A 96-well magnet (Invitrogen, 12027) was used for all subsequent steps and each wash was 175 µl. Samples were washed twice with RIPA buffer, two times with RIPA-500 buffer (10 mM Tris-HCl pH 8, 500 mM NaCl, 1 mM EDTA, 0.1% SDS, 1% Triton X-100), twice with LiCl wash buffer (10 mM Tris-HCl pH 8, 1 mM EDTA, 250 mM LiCl, 0.5% NP-40) and once with EB buffer. Beads were resuspended in 40 µl ChIP elution buffer (10 mM Tris-HCl pH 8, 1 mM EDTA, 300 mM NaCl, 0.4% SDS) + 2 µl Proteinase K (NEB, P8107S). ChIP samples were incubated at 55^°^C for 30 min, 68^°^C for 1 h and held at 14^°^C. Reverse crosslinked samples were magnetized in a 96-well magnet and supernatant was transferred to clean well. DNA cleanup was performed by adding 100 µl pre-warmed SPRI beads (1:2 ratio, Beckman Coulter, 082A63881), mixed by pipetting thoroughly and incubated for 5 min. Samples were magnetized for 5 min, supernatant was removed, beads were washed twice with 80% ethanol and then air dried for 5−10 min. DNA was eluted in 20 µl EB buffer and transferred to another well.

#### Library amplification and QC

Libraries were completed and PCR amplified for 13−14 cycles using 25 µl Kapa HiFi HotStart ReadyMix (Roche, 7958927001) and 2 µl 25 µM primer mix containing complementary to Illumina P7 (forward primer) and P5−i5 index−Partial Read1 (reverse primer) sequences. The amplified libraries were purified with 1X SPRI beads volumes and eluted in 25 µl EB buffer. Library concentration was measured using a Qubit fluorometer (Invitrogen, 15723679) and mean library size was determined by a 4150 TapeStation instrument (Agilent Technologies, G2992AA).

### Data analysis

#### Reference genomes

The following genome assemblies and annotations were employed: *Acanthamoeba castellanii* Acastellanii.strNEFF v1 (GCA_000313135.1), *Arabidopsis thaliana* TAIR10.1 (GCA_000001735.2), *Bigelowiella natans* Bigna1 (GCA_000320545.1), *Creolimax fragrantissima* C_fragrantissima_v5 (GCA_002024145.1), *Dictyostelium discoideum* dicty_2.7 (GCA_000004695.1), *Guillardia theta* Guith1 (GCA_000315625.1), *Naegleria gruberi* V1.0 (GCA_000004985.1), *Nematostella vectensis* jaNemVect1.1 (GCA_932526225.1), *Physcomitrium patens* Phypa V3 (GCA_000002425.2), *Saccharomyces cerevisiae* R64-1-1 (GCA_000146045.2), *Spizellomyces punctatus* S_punctatus_V1 (GCA_000182565.2), *Tetrahymena thermophila* JCVI-TTA1-2.2 (GCA_000189635.1).

#### Transposable element annotation

Transposable elements were annotated using EDTA^35^ v2.2 with a provided CDS file. Transposable elements overlapping genes were removed when examining the coverage of histone modifications.

#### iChIP2 data processing

Raw reads files were aligned to the reference genome of their respective species using Bowtie2^50^ v2.5.3 with parameters *-X 2000 --mm*. Low-quality reads (mapping quality < 30) and duplicate reads were removed using Sambamba^51^ v1.0 with parameters *-F “mapping_quality >= 30”* and *markdup -r*, respectively. Resulting BAM files were processed into CPM-scaled Bigwig format using bamCoverage from deepTools^52^ v3.5.4 with parameters -*e --scaleFactor 1 --normalizeUsing CPM --outFileFormat bigwig --binSize 10*. Finally, ChIP-seq peaks were identified using MACS^53^ v2.2.9.1 with parameters *--max-gap 300 -f BAMPE*.

Sample and experiment quality was assessed by computing the FRiP score, defined as the fraction of reads aligning to called peak coordinates. Further quality assessment was performed by comparing Log2(CPM + 1) values distribution (in all 1kb genomic bins with ≥1 read) between control and ChIP-seq experiments.

Replicates deemed to be of sufficient quality were merged using samtools^54^ v1.21. Bigwig and bedgraph coverage files were generated using bamCoverage with 10bp windows, extending paired end reads, and coverage normalized to the effective genome size, and further log2 normalized to input using bigwigCompare, where both tools are part of deepTools^52^ v3.5.1. Narrow peaks were called for histone modifications and broad peaks were called for H3 and input samples using MACS^53^ v3. Broad peaks were later used to mask regions for histone modification coverage analyses. Genomic data were imported and manipulated in R v4.1.2 using the packages tidyverse^55^ v1.3.1, rtracklayer v1.54.0, reshape2^56^ v1.4.4, dplyr v1.1.4, tidyr v1.3.1, valr^57^ v0.6.6, GenomicRanges^58^ v1.46.1 and plotted using ggplot2 v3.5.1.

#### RNA-seq data sources and processing

Gene expression data were downloaded from the NCBI Sequence Read Archive (SRA) and converted into the fastq format using SRA Toolkit v2.9.0, and then mapped to their corresponding genome using STAR^59^ v2.7.10a. The SRA accession numbers are the following: *Acanthamoeba castellanii* (sample SRR11193414; BioProject PRJNA487265), *Arabidopsis thaliana* (samples XXX; BioProject XXX), *Bigelowiella natans* (samples SRR8306016, SRR8306018, SRR8306019; BioProject PRJNA509798), *Creolimax fragrantissima* (samples SRR2011335, SRR2011336, SRR2011337; BioProject PRJNA283212), *Dictyostelium discoideum* (samples XXX; BioProject XXX), *Guillardia theta* (sample SRR1294409; BioProject PRJNA248394), *Naegleria gruberi* (sample ERR2215457; BioProject PRJEB23822), *Nematostella vectensis* (samples SRR987318−28; BioProject PRJNA213177), *Physcomitrium patens* (samples XXX; BioProject XXX), *Saccharomyces cerevisiae* (samples SRR17310294, SRR17310295; BioProject PRJNA791636), *Spizellomyces punctatus* (samples SRR343039, SRR343043; BioProject PRJNA37881), *Tetrahymena thermophila* (samples SRR13245153, SRR13245156, SRR13245157; BioProject PRJNA684669).

#### Chromatin states

Chromatin segmentation was performed using ChromHMM^30^ v1.24 by first binarizing merged bam files into 200bp windows using BinarizeBam and second using LearnModel with default settings across a range of 2-50 chromatin states. To determine the number of chromatin states to set per species, sequential chromatin state models were correlated in terms of emission probabilities and feature overlap enrichment between sequential chromatin state models. When the correlation of either neared 1 for the lowest number of chromatin states, these were manually inspected to select a low number of unique chromatin states, either in terms of the emission probabilities or overlap with genomic features.

To integrate chromatin states across species, we performed a Pearson correlation among all chromatin states from all species and hierarchically clustered the distances of the correlated states. We manually removed null states prior to hierarchical clustering, which are those chromatin states lacking emission probabilities greater than 0.5 for any histone modification. To group similar chromatin states and simplify the resulting tree, we cut the tree at a height of 0.55, a value manually determined to minimize the number of merged clusters, termed metastates, while maintaining the diversity of chromatin states. The emission probabilities of the resulting metastates are the median values of the individual chromatin state emission probabilities. The percentage of the genome covered by a metastate within a species was calculated by summing the percentage of the genome covered by each chromatin state within the corresponding metastate from the relevant species.

#### Genome browser snapshots

Genome browser snapshots were taken using the package Gviz^60^ v1.38.4 in R v4.1.2. Gene annotation files and histone modification coverage files were imported using valr^57^ v0.6.6. Gene orientation was manually added after plotting.

#### hPTM coverage analysis

Log2-normalized histone modification profiles were clustered using k-means clustering over genes using deepTools^52^ v3.5.1. Coverage files were binned into 10bp bins and only regions between the transcription start site (TSS) and transcription end site of genes were used for clusting into six (or 8 for *Guillardia theta*) clusters. Clusters were reordered from highest to lowest based on average gene expression. Clusters were replotted to include 500bp upstream and 3kb downstream of the TSS for both heatmaps and profile plots (**Fig. 2a, Extended Data Fig. 4**) or replotted to include 5kb upstream and downstream, removing any genes within a cluster that had another gene TSS within 5kb of its TSS to remove confounding effects (**Extended Data Fig. 4**). Unannotated genes were indicated based on the presence of RNA-seq reads and histone modifications typical of TSSs. The presence of reads spanning splice junctions further supported the hypothesis that these were unannotated genes and not spurious antisense transcripts. As with genes, transposable elements were clustered based on 10bp binned coverage of histone modifications, except a variable number of k-means clusters were used, manually determined to represent various types of chromatin found over transposable elements. Clusters were reordered to group together visually similar clusters. Replotted elements included 500bp upstream and downstream.

To calculated histone modification coverage per gene or nucleosome (200bp bin), histone modification coverage per 10bp window was log2 normalized to input and averaged over the feature of interest. For coverage per nucleosome over the gene body per gene cluster, genes were divided into 200bp bins up to 1800bp, plus a 200bp bin from the transcription end site for the last nucleosome and a 500bp bin upstream of the TSS for the promoter. Gene clusters per species are those defined above (**Extended Data Fig. 4)**. Transcript per Million (TPM) values of gene expression were arcsinh-transformed for ease of visualization. Gene lengths per gene cluster were log10-transformed for ease of visualization. For analyses of bimodal TSSs, head-to-head genes were filtered out to avoid confounding signal from upstream gene histone modifications affecting analyses of upstream regions of genes. Genes with a TSS within 5kb were removed from the analysis, as 5kb upstream of the focal gene was plotted. Genes from only one representative gene cluster were plotted.

Mosaic plots were generated in R using the vcd^61^ v1.4-11 package using residual-based shading.

#### Correlation of histone modification coverage and gene expression

Histone modification coverage per gene, log2 normalized to input, was correlated to gene expression with a Spearman correlation per species. To highlight trends in histone modification coverage, genes were divided into five quantiles by gene expression and the histone modification coverage was plotted per quantile.

## Extended Data Figure legends

**Extended Data Figure 1. Summary of antibodies tested in this study.** The table indicates whether the antibodies were successful (green), passed quality checks but with issues such as background noise (blue), or failed entirely in specific organisms (red). Crosses denote the absence of a histone modification in certain species.

**Extended Data Figure 2. Quality control measurements of iChIP2. a**, Barcoding efficiency of 110 ng (light grey) or 440 ng (dark grey) chromatin using different adaptors concentration (1 µM, 2.5 µM or 5 µM). Each condition was tested in triplicates. **b**, Scatter plot shows successful (green), problematic (blue) or failed (red) individual libraries. **c**, Example iChIP2 QC pipeline output. **d**, Examples of successful (left) or unsuccessful (right) reads balancing in different libraries. First and second bar plots indicate the observed versus expected number of reads and the total number of reads per species in a pool for a particular antibody, respectively, while the third bar plot shows the genome size of each individual species. Red, semi-transparent boxes indicate samples that failed to balance.

**Extended Data Figure 3. Hierarchical clustering of ChromHMM chromatin states. a**, Hierarchical clustering of chromatin states calculated separately for each species. Clustering was performed on emission probabilities of each state, where missing states were assigned a value of 0. Boxes around chromatin states indicate those grouped into metastates at a tree height of 0.55. Null states were excluded from the hierarchical clustering. **b**, ChromHMM emission probabilities of each chromatin state from each species. Scale indicated to the right of the plot. **c**, Enrichment of the overlap of each chromatin state with the indicated genomic feature. Enrichment was calculated as ((bp of chromatin state overlapping feature / bp of chromatin state in genome) / (bp of feature in genome / bp of genome)). Scale indicated to the right of the plot. **d**, Null state emission probabilities and overlap enrichment, as in b and c.

**Extended Data Figure 4. Clustering of genes by chromatin modification.** K-means clustering of genes into six to eight clusters per species indicated. Clustering was performed on 10bp bins of histone coverage between the transcription start site (TSS) and transcription end site (TES) and gene clusters are ordered by mean gene expression. Heatmaps show read coverage of a histone modification, log2 normalized to input. Solid vertical lines indicate the TSS and dashed lines indicate the TES, if within 3kb of the TSS. The plotted region includes 500bp upstream of the TSS and 3kb downstream of the TSS. Missing histone modification heatmaps indicate the absence of the histone modification in the species, as determined by mass spectrometry.

**Extended Data Figure 5. Histone modifications and gene expression. a**, Profile plots show average read coverage of a histone modification, log2 normalized to input. Genes are selected from k-means clusters calculated in Extended Data Fig. 4, where genes with the TSS of another gene within 5kb upstream of its TSS were removed. Solid vertical lines indicate the TSS. Solid horizontal lines indicate a value of 0, with values below zero displayed as transparent lines. Profiles extend 5kb before and after the TSS. Browser snapshots of bimodal H3K4me3 around the TSS. Tracks follow the same colour scheme as profile plots, in addition to RNA-seq reads (grey, top) and genes (grey, bottom). Putative unannotated genes are marked with a light grey box in the gene track based on the presence of RNA-seq reads spanning putative splice junctions. **b**, Multiple sequence alignment of Dot1 orthologs from *H. sapiens, S. cerevisiae, A. castellanii, T. brucei*, and *T. cruzi*. The orange star denotes the amino acid position in TbDOT1A important for H3K79 binding and limiting methyltransferase activity to dimethylation. Schematic of *A. castellanii* Dot1 ortholog protein domains are indicated to the right of the alignment. MBD=Methyl binding domain.

**Extended Data Figure 6. Clustering of transposable elements by chromatin modification.** K-means clustering of transposable elements not located within genes. Clustering was performed on histone coverage over the transposable element, with similar clusters ordered together. Heatmaps show read coverage of a histone modificaiton, log2 normalized to input. The plotted region includes 500bp upstream and downstream of the transposable element. Adjacent heatmaps show transposable element classifications by class (left), order (middle), and type (right), where each row corresponds to the displayed transposon in the k-means clustering heatmap. Heterochromatin types summarized in Figure 4a are indicated next to the relevant k-means clusters.

**Extended Data Figure 7. Composition of transposable element clusters by order.** Mosaic plot of transposable element orders and k-means clusters. Colour scale indicates Standardized Pearson residuals, where enrichment is red and depletion is blue, with lighter colours indicating p < 0.05 and darker colours indicating p < 0.0001.

**Extended Data Figure 8. Heterochromatin types across eukaryotes.** Boxplots and browser snapshots illustrating heterochromatin types summarized in Figure 4a. Boxplots show average read coverage of histone modification over a gene, log2 normalized to input. The number of TEs or genes per cluster are indicated above as n‥ Browser snapshot tracks follow the same colour scale as boxplots, in addition to RNA-seq reads (grey, top), transposable elements (black, bottom), and genes (grey, bottom).

