## Extended Data Fig. 1 for "Diversity and evolution of chromatin regulatory states across eukaryotes"

### Extended Data Figure 1

| Antibody | Reference | Host | Clonality | Nvec | Cfra | Scer | Spun | Acas | Ddis | Atha | Ppat | Gthe | Tthe | Bnat | Ngru |
|---|---|---|---|---|---|---|---|---|---|---|---|---|---|---|---|
| | | | | Species | | | | | | | | | | | |
| H3 | ab1791, Abcam | rabbit | polyclonal | Good | Good | Good | Good | Good | Good | Good | Good | Good | Good | Good | Good |
| H3K4me2 | ab32356, Abcam | rabbit | monoclonal | Good | Good | Good | Good | Good | Good | Good | Good | Good | Good | Good | Good |
| H3K4me3 | 07-473, Millipore | rabbit | polyclonal | Good | Good | Good | Good | Good | Good | Good | Good | Good | Good | Good | Good |
| H3K9ac | 17-658, Millipore | rabbit | polyclonal | Good | Good | Good | Good | Good | Good | Good | Good | Good | Good | Good | Good |
| | MA5-33384, ThermoFisher | rabbit | monoclonal | Not profile | Not profile | Not profile | Not profile | Bad | Bad | Bad | Bad | Bad | Not profile | Bad | Bad |
| H3K9me1 | ab9045, Abcam | rabbit | polyclonal | Good | × | × | Good | Good | Good | Good | Good | Pass | × | Pass | × |
| | MA5-33385, ThermoFisher | rabbit | monoclonal | Bad | × | × | Not profile | Bad | Bad | Not profile | Not profile | Not profile | × | Not profile | × |
| H3K9me2 | ab1220, Abcam | mouse | monoclonal | Not profile | Not profile | Not profile | Not profile | Not profile | Not profile | Good | Good | Not profile | Not profile | Not profile | Not profile |
| H3K9me3 | ab176916, Abcam | rabbit | monoclonal | Good | × | × | Good | Good | Good | × | × | × | Pass | Good | Good |
| | C15410056, Diagenode | rabbit | polyclonal | Pass | × | × | Bad | Not profile | Bad | × | × | × | Not profile | Not profile | Not profile |
| | 713008, ThermoFisher | rabbit | polyclonal | Bad | × | × | Not profile | Good | Bad | × | × | × | Not profile | Not profile | Bad |
| H3K27ac | ab4729, Abcam | rabbit | polyclonal | Good | Good | Good | Good | Good | Good | Good | Pass | Good | Good | Good | Pass |
| | 39136, Active Motif | rabbit | polyclonal | Not profile | Not profile | Not profile | Not profile | Not profile | Not profile | Not profile | Not profile | Not profile | Not profile | Not profile | Good |
| H3K27me3 | 07-449, Millipore | rabbit | polyclonal | Bad | × | × | Good | Good | Pass | Good | Pass | Pass | Good | Good | Good |
| | 17-622, Millipore | rabbit | polyclonal | Bad | × | × | Good | Good | Bad | Bad | Not profile | Bad | Bad | Bad | Bad |
| | ab192985, Abcam | rabbit | monoclonal | Bad | × | × | Bad | Bad | Bad | Not profile | Not profile | Bad | Not profile | Not profile | Not profile |
| | 9733, Cell Signaling | rabbit | monoclonal | Bad | × | × | Bad | Not profile | Bad | Not profile | Not profile | Bad | Not profile | Not profile | Not profile |
| | C15410069, Diagenode | rabbit | polyclonal | Bad | × | × | Not profile | Bad | Bad | Not profile | Not profile | Bad | Not profile | Not profile | Bad |
| | MA5-11198, ThermoFisher | rabbit | monoclonal | Good | × | × | Not profile | Bad | Bad | Not profile | Not profile | Bad | Not profile | Bad | Bad |
| H3K36me3 | ab9050, Abcam | rabbit | polyclonal | Good | Good | Good | Good | Good | Good | Good | Good | Good | Good | Pass | Good |
| H3K79me1 | ab177185, Abcam | rabbit | monoclonal | Good | Good | Good | Good | Pass | Good | × | × | Pass | Good | Pass | × |
| H3K79me2 | ab3594, Abcam | rabbit | polyclonal | Good | Good | Good | Good | Good | Good | × | × | × | Good | Pass | × |
| H3K79me3 | C15410068, Diagenode | rabbit | polyclonal | Good | × | Good | Good | Good | Good | × | × | × | Good | Pass | × |
| H4K16ac | 07-329, Millipore | rabbit | polyclonal | Pass | Good | Good | Good | Good | Good | Good | Good | Pass | Good | Good | Good |

● Good ● Pass ● Bad ○ Not profile × Histone modification not present in species
