## Supplementary figures and images for "Diversity and evolution of chromatin regulatory states across eukaryotes"

### Extended Data Fig. 2

#### Extended Data Figure 2

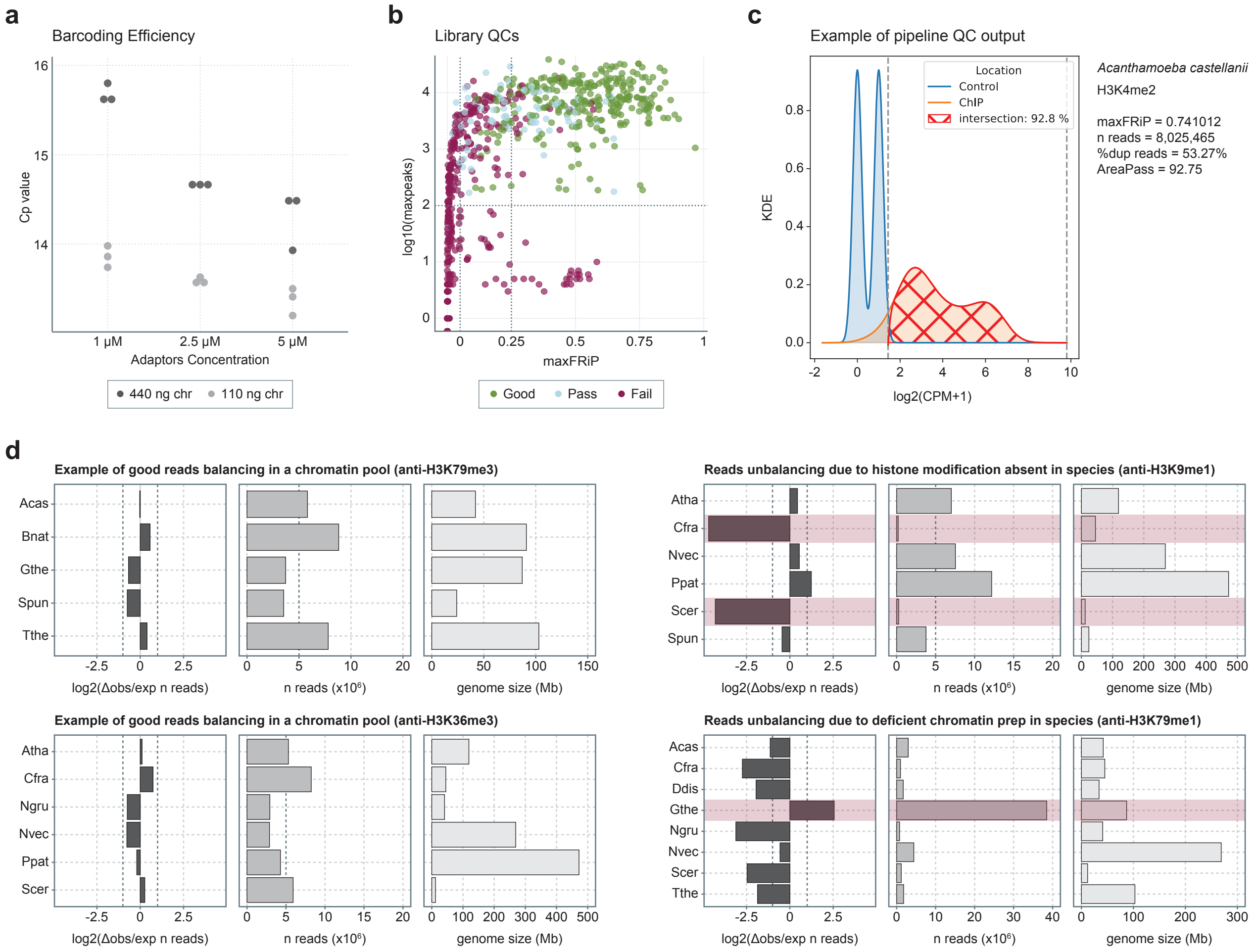

### Extended Data Fig. 3

##### Extended Data Figure 3

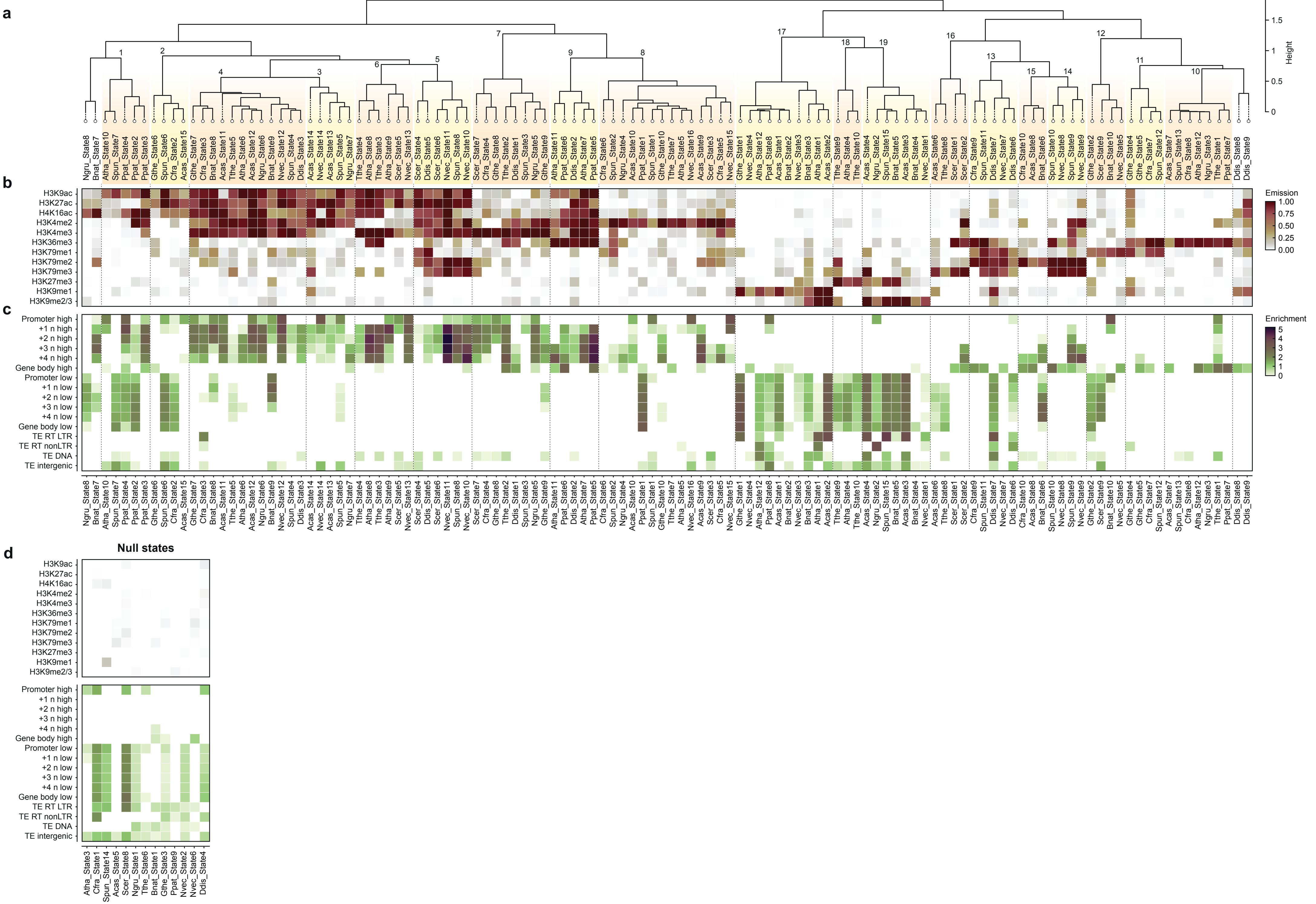

### Extended Data Fig. 4

#### Extended Data Figure 4

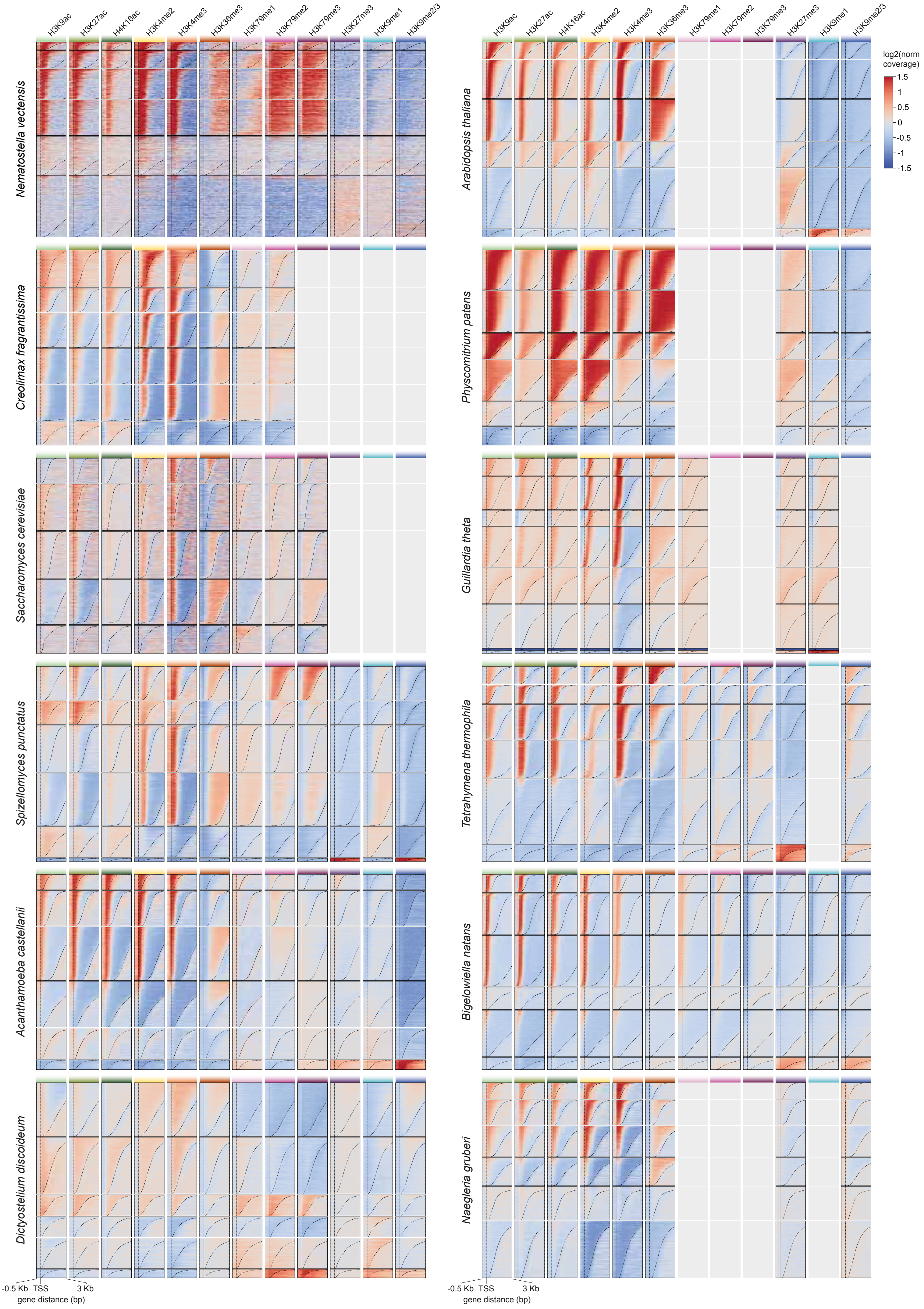

### Extended Data Fig. 5

#### Extended Data Figure 5

**a**

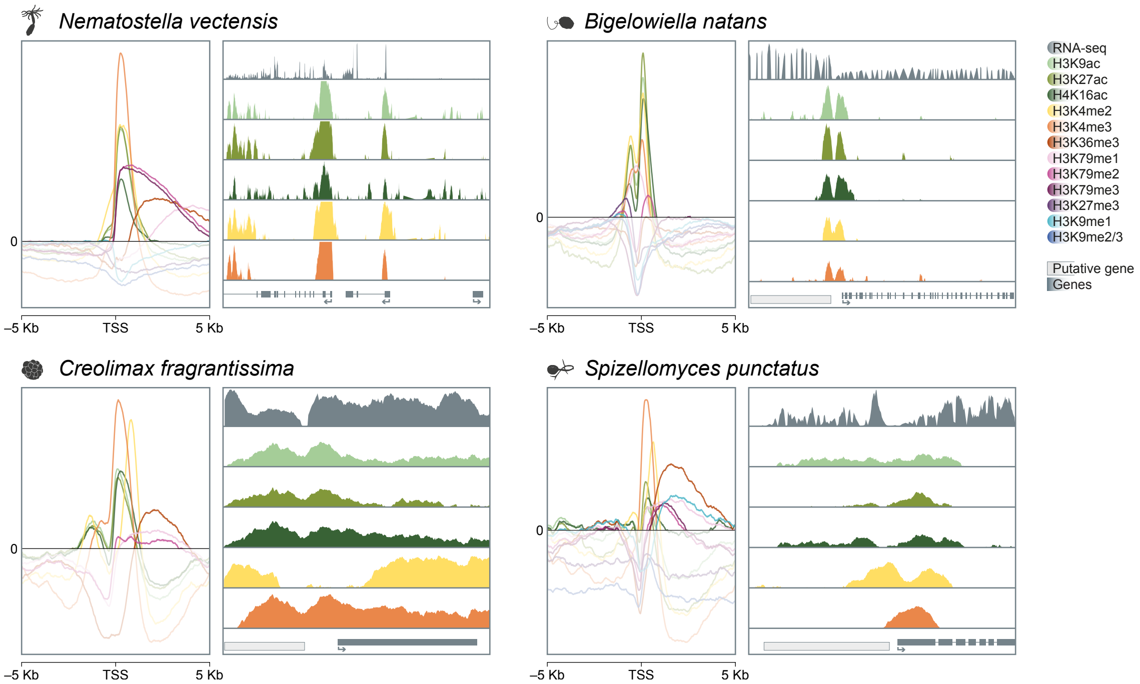


**b**

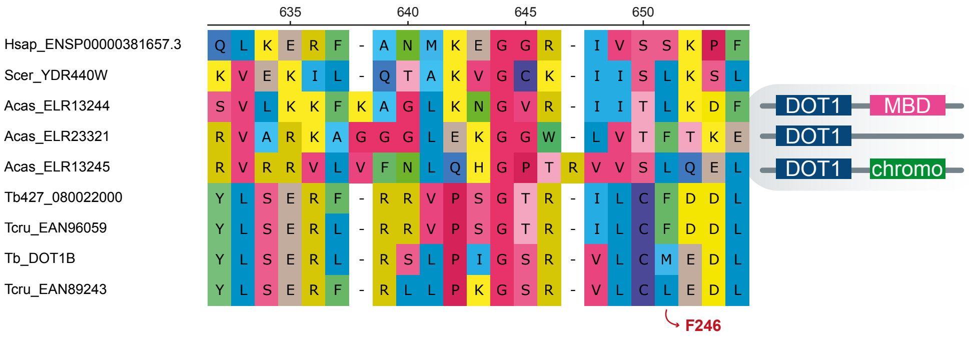

### Extended Data Fig. 6

#### Extended Data Figure 6

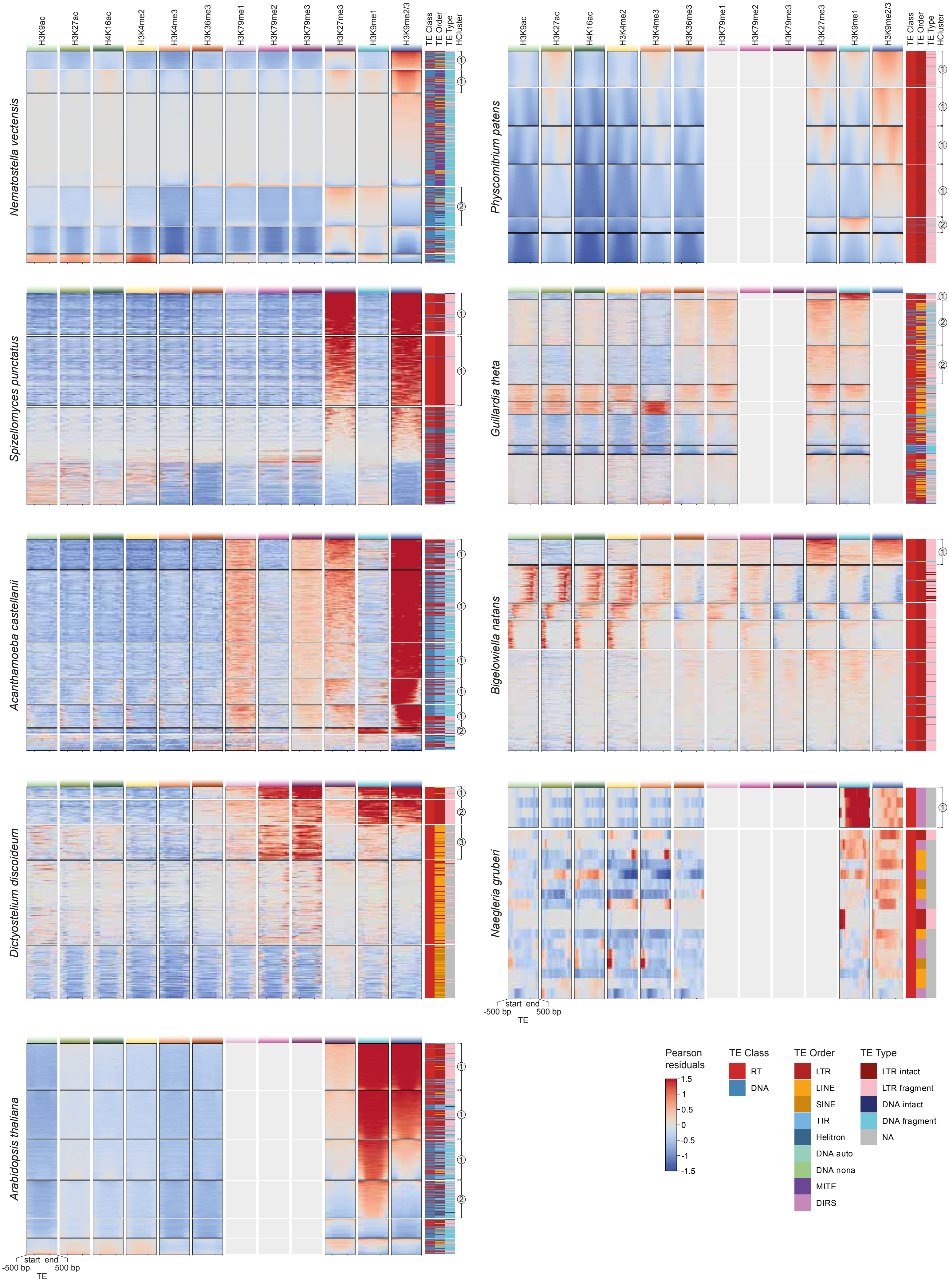

### Extended Data Fig. 7

#### Extended Data Figure 7

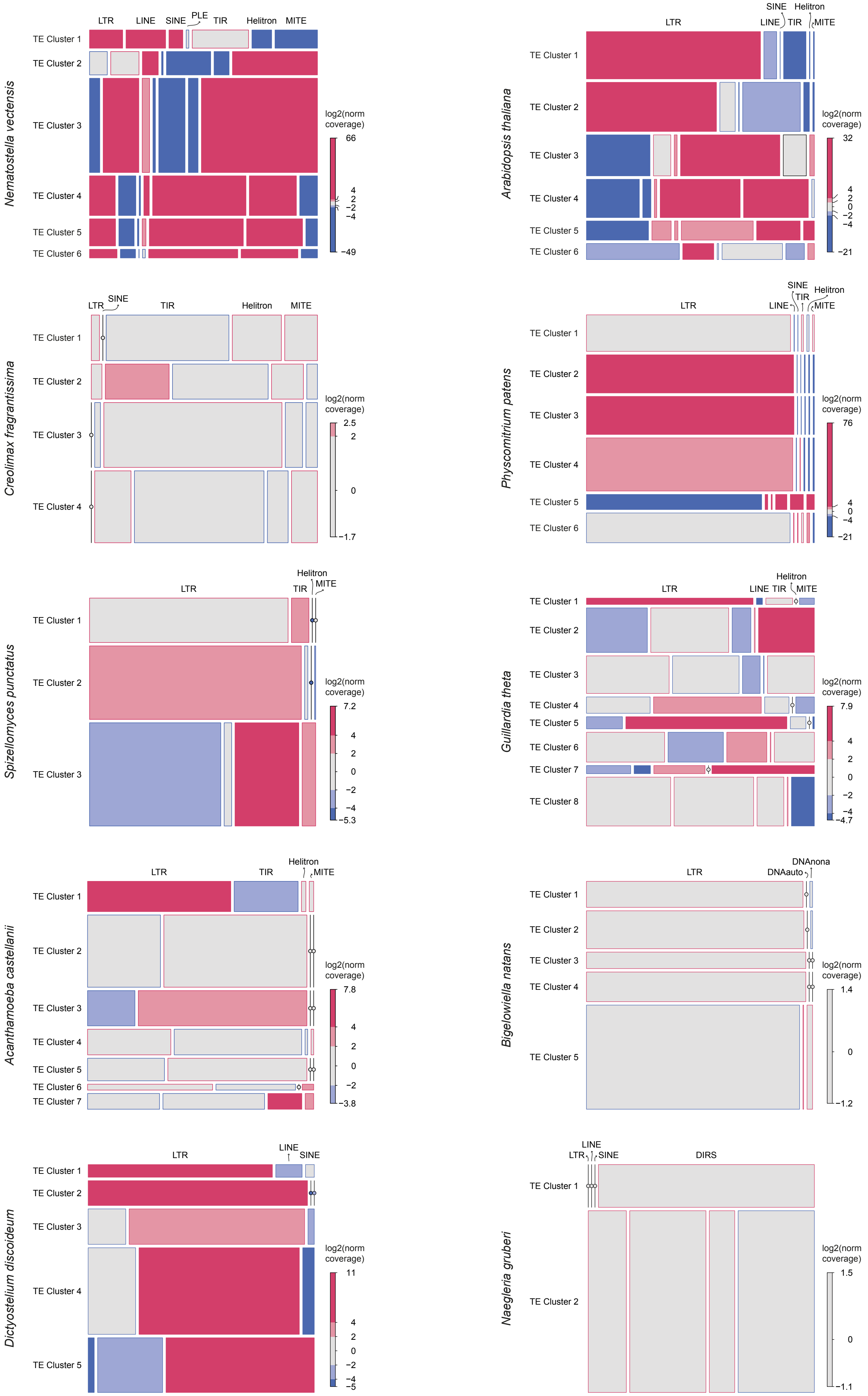

### Extended Data Fig. 8

##### Extended Data Figure 8

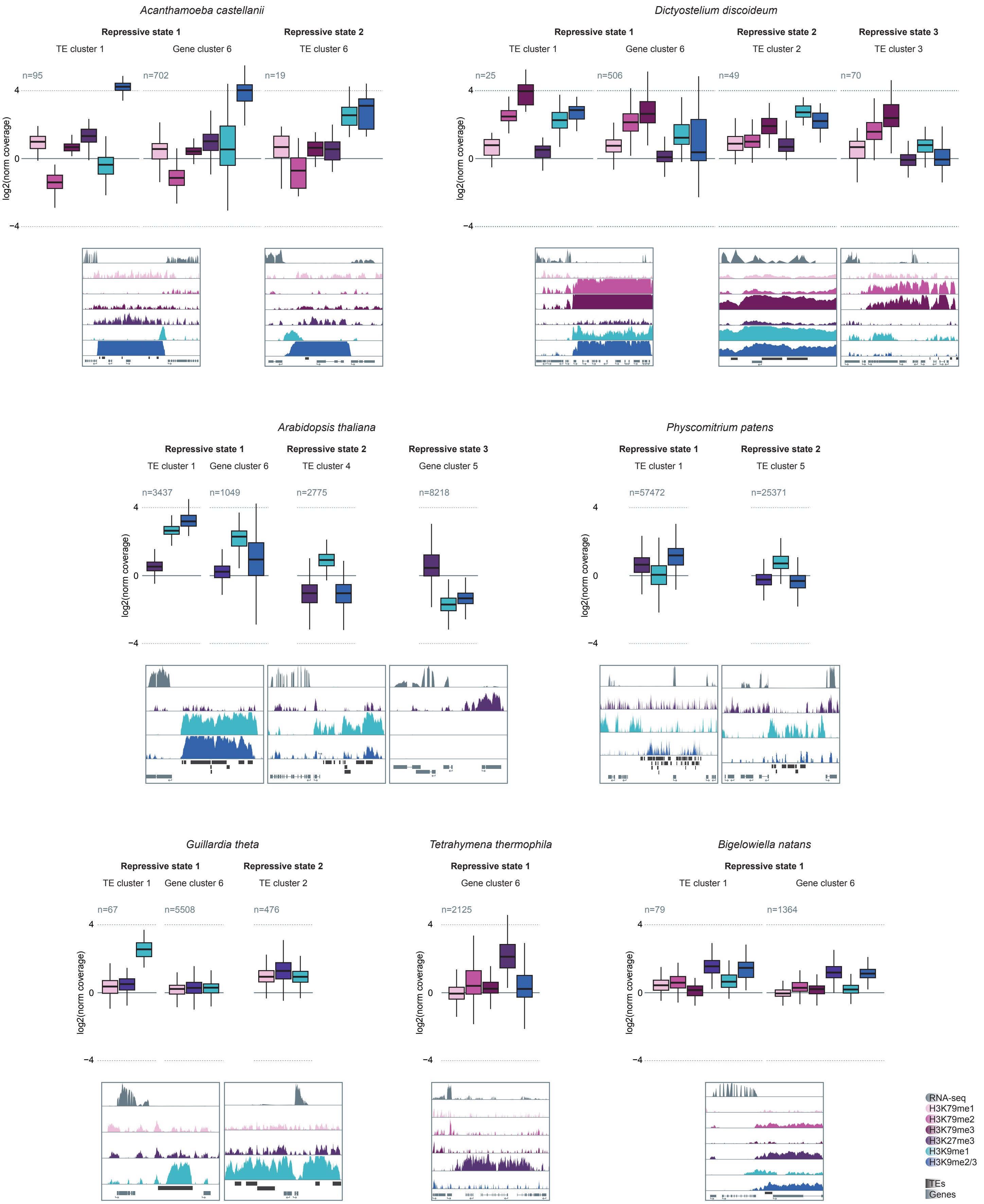
